# Contextual Classifications of Cancer Driver Genes

**DOI:** 10.1101/715508

**Authors:** Pramod Chandrashekar, Navid Ahmadinejad, Junwen Wang, Aleksandar Sekulic, Jan B. Egan, Yan W. Asmann, Carlo Maley, Li Liu

**Affiliations:** College of Health Solutions, Arizona State University, Tempe, AZ, 85004, USA; Biodesign Institute, Arizona State University, Tempe, AZ, 85281, USA; Department of Health Sciences Research & Center for Individualized Medicine, Mayo Clinic Arizona, Scottsdale, AZ, 85259, USA; Department of Health Sciences Research, Mayo Clinic Florida, Jacksonville, AZ, 32224, USA

## Abstract

Functions of cancer driver genes depend on cellular contexts that vary substantially across tissues and organs. Distinguishing oncogenes (OGs) and tumor suppressor genes (TSGs) for each cancer type is critical to identifying clinically actionable targets. However, current resources for context-aware classifications of cancer drivers are limited. In this study, we show that the direction and magnitude of somatic selection of missense and truncating mutations of a gene are suggestive of its contextual activities. By integrating these features with ratiometric and conservation measures, we developed a computational method to categorize OGs and TSGs using exome sequencing data. This new method, named genes under selection in tumors (GUST) shows an overall accuracy of 0.94 when tested on manually curated benchmarks. Application of GUST to 10,172 tumor exomes of 33 cancer types identified 98 OGs and 179 TSGs, >70% of which promote tumorigenesis in only one cancer type. In broad-spectrum drivers shared across multiple cancer types, we found heterogeneous mutational hotspots modifying distinct functional domains, implicating the synchrony of convergent and divergent disease mechanisms. We further discovered two novel OGs and 28 novel TSGs with high confidence. The GUST program is available at https://github.com/liliulab/gust. A database with pre-computed classifications is available at https://liliulab.shinyapps.io/gust

## INTRODUCTION

In tumor development and progression, two groups of genes work synergistically to promote and maintain abnormal cell growth (1,2). One group is oncogenes (OGs) that cause cancers via gain of functions. The other group is tumor suppressor genes (TSGs) that cause cancers via loss of functions. Activities of these genes depend strongly on cellular contexts. Except a few cases such as *TP53* and *RAS* that are common drivers of many different cancer types, most OGs and TSGs have high tissue specificities and behave as passenger genes (PGs) outside the contexts (3–5). Some genes even play dual roles. An evident example is the *NOTCH1* gene that is an OG in hematologic malignancies such as acute lymphoblastic leukemia, but is a TSG in solid tumors such as lung cancers, melanoma and hepatocellular carcinoma (6–8). Consistent with the contextual activities, therapies targeting the Notch signalling pathway have highly variable efficacy and have been associated with acquired drug resistance (9). Thus, identification of driver genes and further distinguishing OGs and TSGs in the context of individual cancer types are critical to understanding cancer etiology and developing effective treatments.

While there are abundant resources for classifying cancer drivers from passengers, very few recognize contextual distinctions. The widely used cancer gene consensus (CGC, version 87, last accessed in February 2019) (10) curates 723 confirmed or putative cancer drivers, among which less than half are categorized as OGs or TSGs. Another collection of manually curated cancer drivers from OncoKB (last accessed in April 2019) (11) documents 256 OGs and 271 TSGs, among which only 52 genes have information related to cancer types. Furthermore, these annotations are mainly for well-studied cancers e.g., melanoma and colorectal cancers. For other cancers, e.g., cholangiocarcinoma and thymoma, CGC and OncoKB contain no information even though genomic profiles of these tumors are readily available.(12,13)

Complementary to time-consuming manual curations and experimental assays, computational predictions can quickly prioritize potential cancer drivers using high-throughput omics data. A recent study surveyed 26 computational tools for cancer driver predictions.(14) However, one only method (20/20+) contrasts OGs and TSGs.(15) This method is an extension of the conventional 20/20 rule that requires an OG to have >20% mutations causing missense changes at recurrent positions, and requires a TSG to have >20% mutations causing inactivating changes.(16) However, this rule does not extend beyond a small set of canonical examples. Because many TSGs harbor hotspots of missense mutations (17,18), recurrent missense mutations are not a deterministic feature of OGs. Meanwhile, a stochastic mutational process can introduce inactivating mutations into OGs and PGs that later increase in frequency via genetic drift (19–21). Therefore, conventional ratiometric measures such as mutational frequencies and clustering patterns need to be augmented with additional informative features.

Because tumor development is an evolutionary process, characteristics of the mutation-selection interplay can be instructive. For example, a mutational rate higher than background is a strong indicator of driver genes that has been encompassed in several tools (22,23) although this feature does not distinguish OGs and TSGs. To overcome this limitation, we investigated a new set of features that quantify the direction and magnitude of selection of cancer genotypes. In an evolutionary framework, positive selection promotes advantageous genotypes in favor of a higher fitness of a tumor; negative selection eliminates genotypes with adverse effects; and neutral selection allows insignificant genotypes to drift (24–26). Because an OG promotes tumor growth via missense mutations causing gain of functions, these activating mutations are positively selected, and inactivating mutations are negatively selected. Conversely, because a TSG is disabled in tumors, protein-truncating mutations are positively selected and other mutations are under neutral selection. For passenger genes (PGs), all mutations are under neutral selection (25,27). When activities of a gene vary significantly across cancer types, the direction and magnitude of somatic selection will change accordingly, enabling contextual classification of driver genes.

Under this premise, we developed a computational method, named genes under selection in tumors (GUST) that integrates somatic selection of genes in tumor development, molecular conservation during species evolution and conventional ratiometric measures to classify OGs, TSGs and PGs. We applied this method to categorizing driver genes in 33 cancer types from the cancer genome atlas (TCGA) (28). Our analysis reveals excessive context-specific drivers. We conclude with a discussion on the clinical implications of these findings.

## MATERIAL AND METHODS

GUST is a random forest model that predicts the class label (OG, TSG or PG) of a gene based on 10 meticulously engineered features from exome sequencing data. In the following sections, we first describe the compilation of training data, then the construction of various features, and finally the configuration of a multiclass random forest classifier.

### Training gene set

Because existing cancer gene databases (e.g., OncoKB and CGC) do not provide context-dependent annotations, current methods are trained on data aggregated from multiple cancer types to make predictions. Since many driver genes have tissue-specific functions (14), this strategy risks introducing biases into the models. To mitigate this problem, we compiled the first-ever collection of OGs, TSGs and PGs in a cancer-type specific manner. We began with two lists of genes with complementary information. The first list consisted of 36 OGs, 48 TSGs and 21 genes with dual OG/TSG roles annotated in the CGC. The tumor activating or suppressing roles of these genes have been confirmed with cancer hallmarks in experimental assays and are attributable to coding substitutions or indels (29). The second list consisted of 235 computationally predicted driver genes that were assigned to specific cancer types (14). These predictions were based on a meta-analysis of the TCGA samples with 26 computational programs. These two lists shared 70 genes. We then retrieved somatic mutations of these 70 genes in 10,172 tumor samples representing 33 cancer type from the TCGA project (28). For a gene to qualify as an OG in a specific cancer type, it needs to be annotated as an OG or a dual-role gene in the CGC, predicted as a driver in the meta-analysis of the matching cancer type, and display mutational hotspots in the corresponding TCGA tumor samples. For a gene to qualify as a TSG in a specific cancer type, it needs to be annotated as a TSG or a dual-role gene in the CGC, predicted as a driver in the meta-analysis, and have an overabundance of truncating mutations in the corresponding TCGA tumor samples. For a gene to qualify as a PG in a specific cancer type, it needs to be predicted as a PG in the meta-analysis and shows no mutational hotspots nor overabundance of truncating mutations in corresponding TCGA tumor samples. Genes that did not meet these requirements were removed. The final collection consisted of 55 OG annotations, 174 TSG annotations and 304 PG annotations that involved a total of 50 known driver genes and 33 cancer types (**Supplementary Table 1**).

### Somatic selection features

Given a gene with somatic mutations reported in a collection of tumor samples, we denote the selection coefficient of missense mutations as ω, and the selection coefficient of protein-truncating (nonsense and frameshifting) mutations as φ. To account for different mutational rates, we consider seven substitution types (1: A→C or T→G, 2: A→G or T→C, 3: A→T or T→A, 4: C→A or G→T, 5: C→G or G→C, 6: C→T or G→A at non-CpG sites, and 7: C→T or G→A at CpG sites), one insertion type and one deletion type. Based on the statistical framework proposed by Greenman *et. al*,(30) the probability of observing these mutations is a product of multinomial distributions

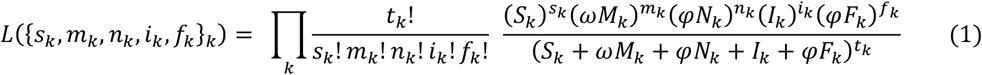

where *s*_*k*_, *m*_*k*_, *n*_*k*_, *i*_*k*_ and *f*_*k*_ are the observed numbers of synonymous, missense, nonsense, in-frame and frameshifting mutations in the *k*^*th*^ rate category, respectively, *S*_*k*_, *M*_*k*_, *N*_*k*_, *I*_*k*_ and *F*_*k*_ are the corresponding expected numbers of changes computed by saturated mutations, and *t*_*k*_ = *s*_*k*_ + *m*_*k*_ + *n*_*k*_ + *i*_*k*_ + *f*_*k*_ is total number of observed mutations. The values of log(ω) and log(φ) are determined by maximizing the log likelihood *L* and constrained within the range of [−5, 5]. The sign and absolute value of log(ω) and log(φ) indicate the direction and magnitude of somatic selection. Values around 0 indicate neutral somatic selection.

### Ratiometric and conservational features

For each gene, we computed traditional ratiometric parameters measuring the fraction of missense mutations and the fraction of truncating mutations among all mutations. To detect mutational hotspots, we first applied density estimates with a rectangular kernel and a bandwidth of 5 base pairs to aggregate closely-spaced missense mutations into peaks. We then calculated the fraction of mutations inside peaks and the fraction of mutations inside the highest peak. For truncating mutations, we computed the average ratio of lengths of the truncated peptides relative to the length of the wild type form. To estimate evolutionary conservation of a gene, we retrieved multiple sequence alignments of 100 vertebrate species from the UCSC Genome Browser database, and computed the substitution rate of each position using the Fitch algorithm (31). The average substitution rate over all positions measures gene-level conservation. The average substitution rate over positions in a summit measures conservation of the mutational hotspot. Altogether, the GUST method uses 10 features (**Table 1**).

**Table 1.** The ten predictors used by GUST

| Category | Predictor | Description |
| --- | --- | --- |
| Somatic selection | $\log(\omega)$ | Selection coefficient of missense mutations |
| | $\log(\phi)$ | Selection coefficient of truncating mutations |
| Ratiometrics | R.missense | Fraction of missense mutations over all mutations |
|  | R.truncating | Fraction of truncating mutations over all mutations |
|  | R.peak | Fraction of missense mutations under peaks |
|  | R.summit | Fraction of missense mutations under the highest peak |
|  | C.summit | Count of missense mutations under the highest peak |
|  | R.length | Mean fraction of lengths of truncated proteins |
| Conservation | E.gene | Mean substitution rate of all positions of the protein |
|  | E.summit | Mean substitution rate of positions under the highest peak |

### Random forest classifier

Each record in the training data involved one gene and one cancer type. For a given gene/cancer-type pair, we retrieved somatic mutations from the corresponding TCGA tumor samples and computed values of the 10 features. Using these features as predictors and the class labels as the response, we constructed a random forest classifier with 200 trees. For each gene, this model produces three probability scores of it being an OG, a TSG or a PG, respectively. It assigns the class label based on the highest probability score. To estimate the sensitivity and specificity associated with a prediction, we built ROC curves for one-vs-rest predictions and extrapolated the values corresponding to the probability score.

### Preprocessing TCGA data

The TCGA project provides whole-exome sequencing data of 10,172 tumor samples representing 33 cancer types(28). We retrieved somatic mutations called by the GATK/MuTect pipeline (32) against the hg38 human reference genome from the Genomic Data Commons (GDC) data portal (33). We removed low-quality mutations and kept single nucleotide substitutions causing synonymous, missense or nonsense changes, and indels causing in-frame or frame-shifting changes of the encoded proteins. Due to the rare occurrences (<1%) and low confidence (34) of predicted splice site mutations, we did not include these mutations in the analysis. We computed the mutational load of a tumor as the number of mutations it contained. For each cancer type, we removed samples with mutational loads outside the 1.5 interquartile range below the first quartile or above the third quartile, respectively. A total of 194 hypo-mutated and 315 hyper-mutated were removed. For each cancer type, we removed less frequently mutated genes that had fewer than 4 protein-altering mutations or were mutated in less than 2% of tumor samples. Because uterine corpus endometrial carcinoma had a significantly higher mutational load than the other cancer types (z test p-value=0.017), we increased the threshold for this cancer type to remove genes mutated in less than 5% of tumor samples.

## RESULTS

### Evaluation of the GUST method

#### Prediction accuracy

We split the training genes into ten equal-sized disjoint subsets and evaluated the random forest model via 10-fold cross validations. The testing accuracy of this model was 0.94. For comparison, we searched for existing tools that predict OGs and TSGs. Among the three tools we found, only 20/20+ (15) is actively maintained. Because executables of the other two methods (35,36) are not available, we did not include them in the evaluation. From the GDC data portal (33), we retrieved precomputed 20/20+ predictions of the gene/cancer-type pair in the training set. The overall accuracy of 20/20+ was 0.86, 8% lower than GUST. To calculate traditional performance metrics, we converted three-class predictions to binary predictions by contrasting one class with the other two classes combined, i.e., one-vs-rest predictions. In all categories, GUST showed better or comparable performance than 20/20+. The largest improvements were on precisions of identifying OGs or TSGs, which increased from 0.78 – 0.82 in 20/20+ to 0.91 – 0.94 in GUST, showing up to 15% improvement (**Table 2**).

**Table 2.** Performance of GUST and 20/20+

| Binary Classes |  | GUST |  |  |  |  |  | 20/20+ |  |  |  |  |  |
| --- | --- | --- | --- | --- | --- | --- | --- | --- | --- | --- | --- | --- | --- |
| Positive | Negative | TPR | TNR | PPV | NPV | Acc. | AUC | TPR | TNR | PPV | NPV | Acc. | AUC |
| OG, TSG | PG | 0.94 | 0.95 | 0.94 | 0.95 | 0.95 | 0.98 | 0.90 | 0.85 | 0.82 | 0.92 | 0.88 | 0.94 |
| OG | PG, TSG | 0.89 | 0.99 | 0.92 | 0.99 | 0.98 | 0.99 | 0.95 | 0.97 | 0.78 | 0.99 | 0.97 | 0.97 |
| TSG | PG, OG | 0.93 | 0.96 | 0.91 | 0.96 | 0.95 | 0.98 | 0.86 | 0.90 | 0.81 | 0.93 | 0.89 | 0.93 |
| Three Classes |  | - | - | - | - | 0.94 | 0.98* | - | - | - | - | 0.86 | 0.95* |

TPR: true positive rate, sensitivity; TNR: true negative rate, specificity; PPV: positive predictive value, precision; NPV: negative predictive value; Acc.: Accuracy; AUC: Area under the ROC curve.
\* Macro-AUC values were calculated by averaging three one-vs-rest ROC curves. Linear interpolation was used between points of ROC (60).

The ROC curves for one-vs-rest predictions reconfirmed the superior performance of GUST (**Fig. 1A**). Compared to 20/20+, GUST had a significantly higher AUC value of the PG-vs-rest ROC curve (0.98 vs. 0.94, DeLang’s test p-value=0.0008), and a significantly higher AUC value of the TSG-vs-rest ROC curve (0.98 vs. 0.93, p-value=0.001). However, the AUC values of the OG-vs-rest ROC curves were not significantly different between these two methods (0.99 vs. 0.97, p-value=0.21).

**Figure 1.**
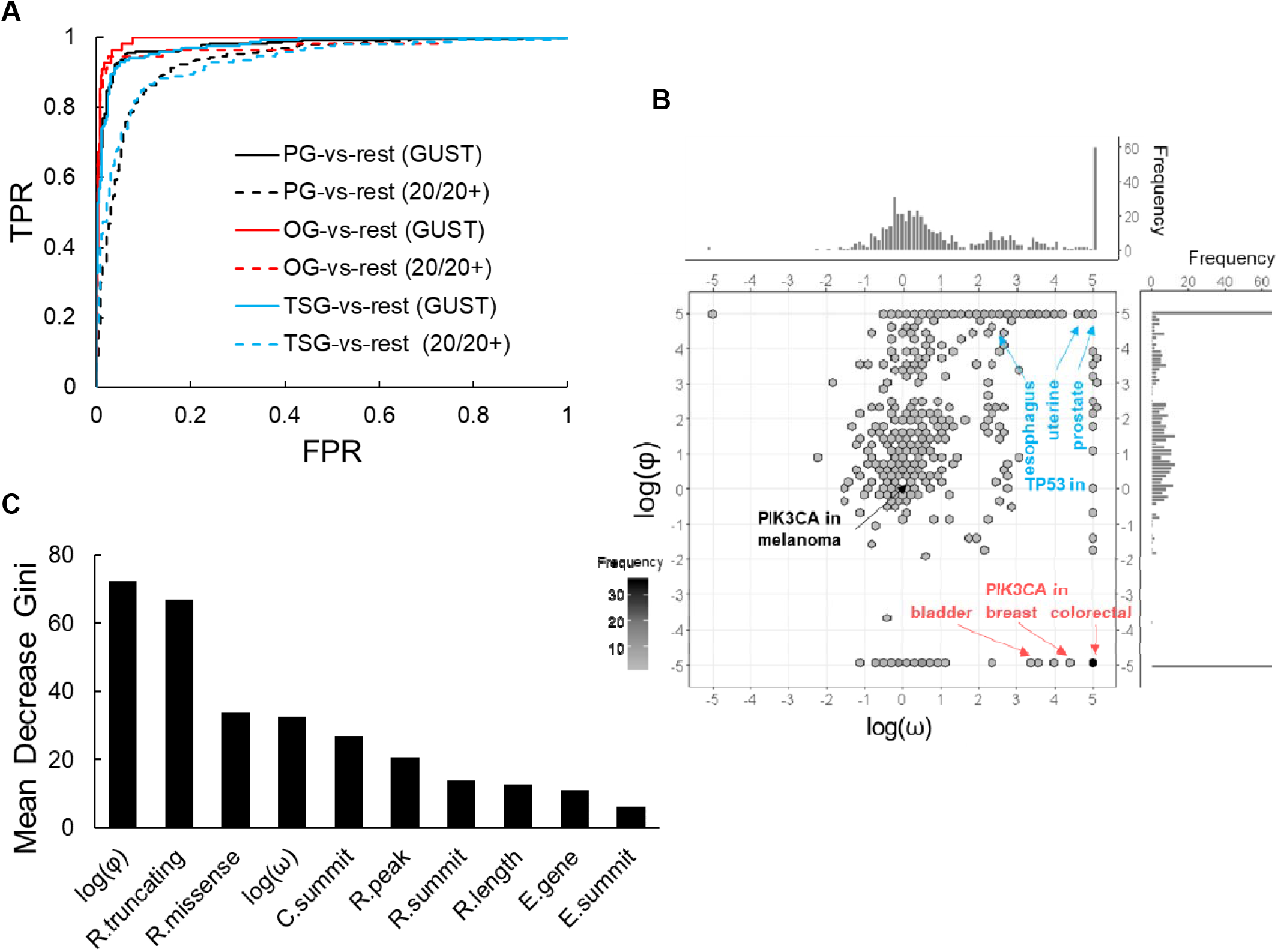
The GUST method. (**A**) ROC curves of one-vs-rest predictions for GUST and for 20/20+. (**B**) Distributions of log(ω) and log(φ) values of 50 genes in 33 cancer-type in the training set. Shades of hexagon bins represent the number of observations. (**C**) Variable importance of each feature in the random forest model.

#### Model interpretation

We first examined the somatic selection features. As expected, well-known OGs such as *PIK3CA* in bladder cancers (37), breast cancers (38) and colorectal cancers (39) had high log(ω) values and low log(φ) values (**Fig. 1B**). TSGs such as *TP53* in esophageal cancers (40) uterine cancers (41) and prostate cancers (42) had high log(φ) values. When a gene switches from a driver’s role in one cancer type to a passenger’s role in another cancer type, the functional change is reflected on selection coefficients. For example, the log(ω) and log(φ) values of *PIK3CA* converted to close to zero in melanoma indicating neutral selection. Indeed, the passenger role of *PIK3CA* in melanoma has been proposed in a recent study that shows *PIK3CA*-mutated melanoma cells rely on cooperative signaling to promote cell proliferation and PI3K inhibitors do not repress tumor growth in the absence of other activating driver genes in melanoma (43).

To measure the importance of each predictor in the random forest model, we computed the mean decreased Gini index by permuting out-of-bag data (44). The most informative predictors are the selection coefficient and fraction of truncating mutations, followed by the selection coefficient and fraction of missense mutations (**Fig. 1C**). The combined variable importance of the remaining ratiometric features was lower than that of log(φ). Sequence conservation features had the lowest importance, which was in stark contrast to models predicting pathogenic mutations in germline cells (45).

### Application to TCGA data

During model development, we used a small subset of TCGA data to compute feature values for the training genes. Here, we applied the GUST method to the complete collection of TCGA samples to perform context-aware gene classifications. In total, we identified 98 OGs and 179 TSGs that potentially promoted tumorigenesis for at least one cancer type (**Fig. 2A**, **Supplementary table 2-3**). Among these putative driver genes, 77.6% OGs and 40.2% TSGs were rare with mutations detected in less than 10% of samples of a specific cancer type. The number of rare drivers identified by GUST was positively correlated with mutational load and sample size (multiple linear regression p-values<10^−3^, **Fig. 2B**). The number of common drivers increased only with mutational load (p-value<10^−4^) but not with sample size (p-value=0.72, **Fig. 2C**).

**Figure 2.**
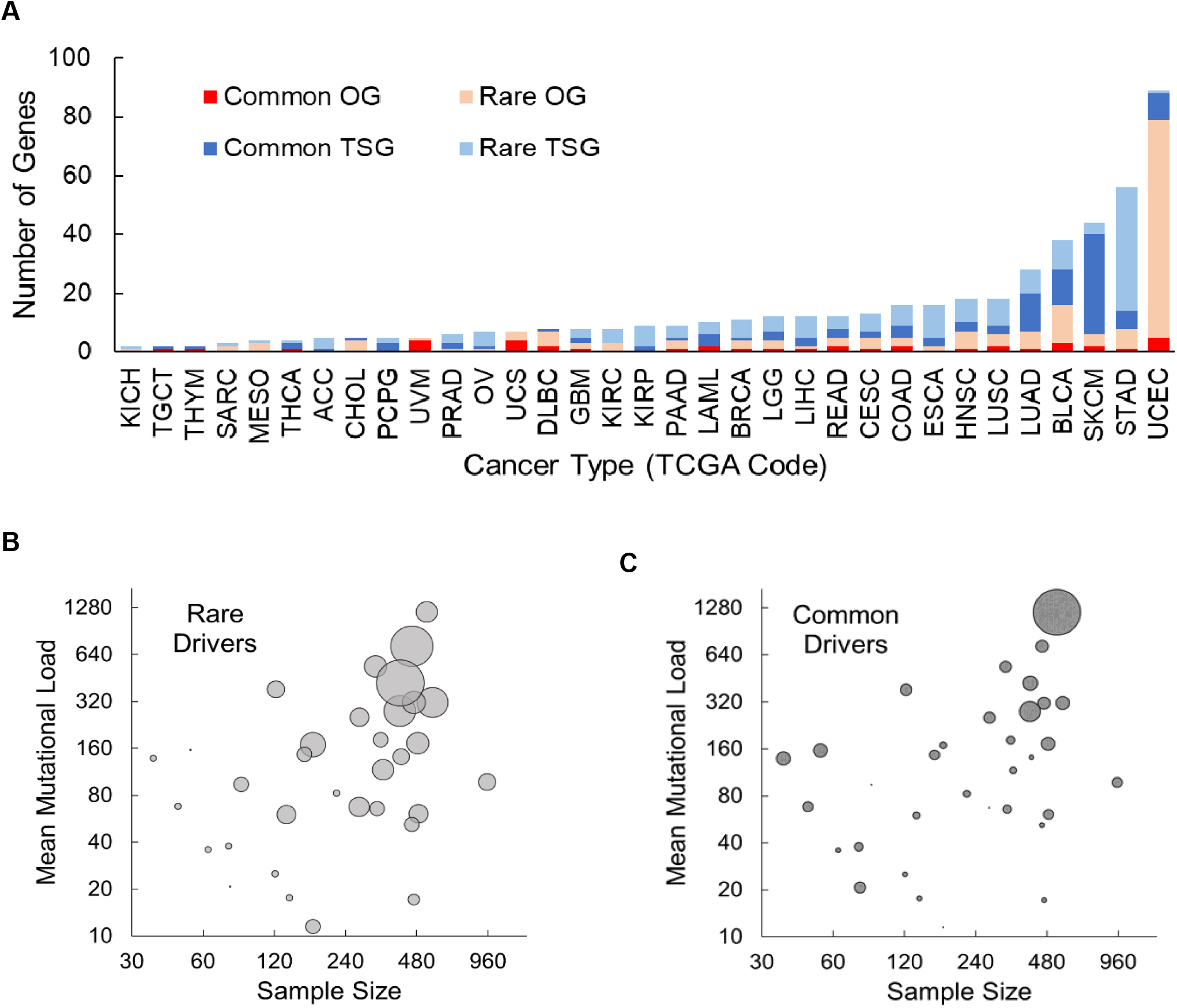
GUST analysis of the TCGA samples. (**A**) Number of common and rare OGs and TSGs found in each cancer type. Abbreviations of cancer types are listed in Supplementary Table 2. (**B, C**) Bubble plot showing the relationship among sample size, mean mutational load and the number of drivers found in each cancer type. Size of bubbles is proportional to the number of rare drivers (**B**) and the number of common drivers (**C**).

#### Spectrum of tissue specificities

Most of these putative drivers were engaged in only one cancer type, showing high tissue specificities. Broad-spectrum drivers that promoted tumorigenesis in two or more cancer types accounted for only 17.3% OGs and 28.5% TSGs in our analysis (**Fig. 3A**). The most prevalent OG was the *PIK3CA* gene found in 16 cancer types, followed by the *KRAS* gene found in 12 cancer types. The most prevalent TSG was the *TP53* gene found in 23 cancer types, followed by the ARID1A gene found in 15 cancer types. We found that broad-spectrum drivers often displayed heterogeneous mutational profiles. Thirteen out of the 17 broad-spectrum OGs possessed multiple mutational hotspots that were selectively engaged in different cancer types. A representative example was the *EGFR* gene. In lung adenocarcinoma samples, *EGFR* mutations clustered at a single mutational hotspot affecting the tyrosine kinase activation loop. In glioma samples, the mutations clustered at two separate hotspots affecting the extracellular domains independent of kinase activities (**Fig. 3B**). The contextual selection of mutations averting the kinase catalytic domain in glioma suggests alternate adaptation of the *EGFR* signaling pathway. For cancer management, although tyrosine kinase inhibitors blocking *EGFR* are common in the therapeutic armamentarium of lung cancer (46,47), these agents have not been successful in treating glioma even with improved drug delivery techniques to penetrate the blood-brain barrier (48–50). Our findings offer a new direction to investigating and enhancing current treatment regimen.

**Figure 3.**
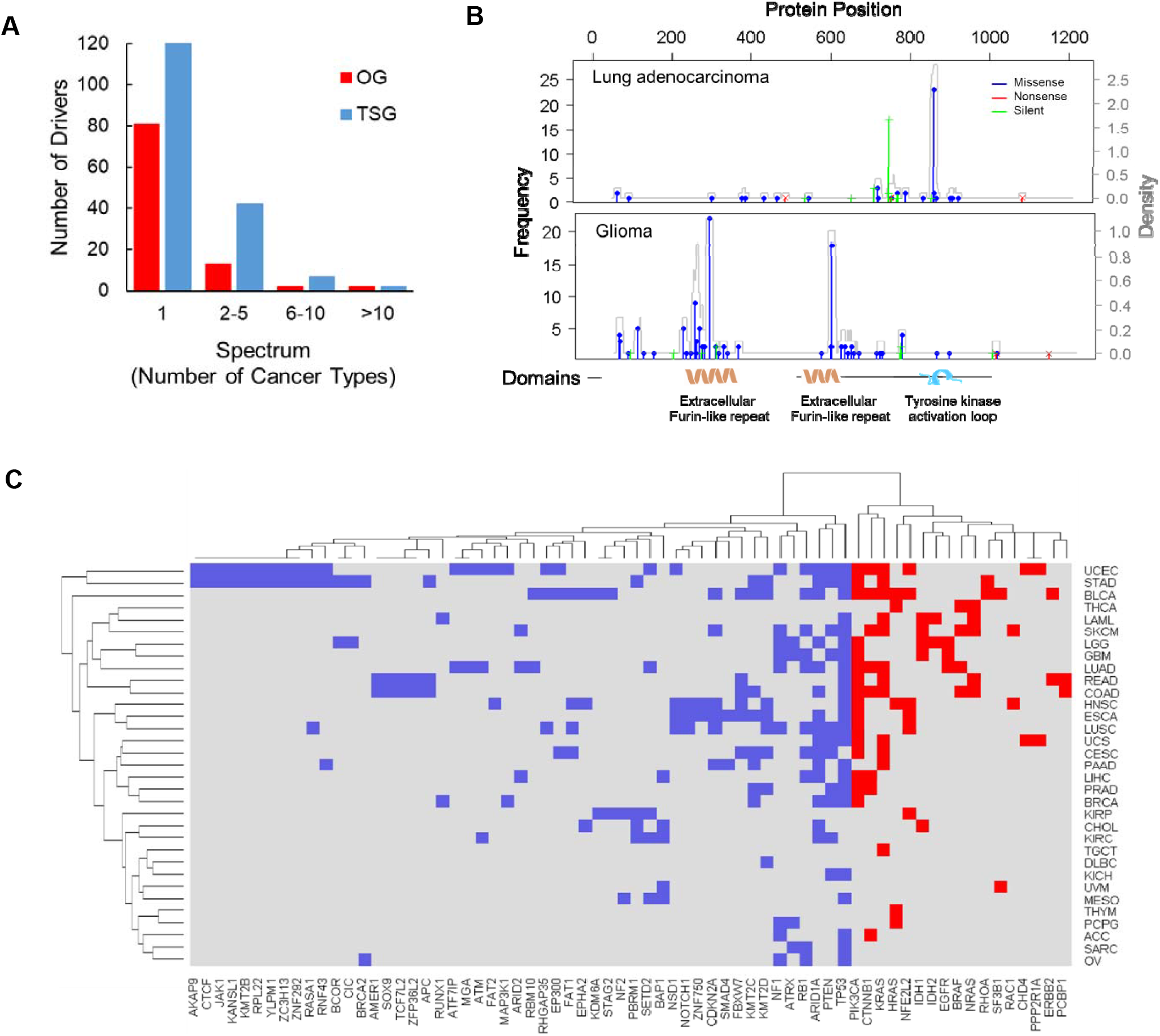
Tissue specificities of cancer drivers. (**A**) Distribution of driver genes with different tissue spectrum. (**B**) Distribution of mutations in the *EGFR* gene in lung adenocarcinoma and glioma (low-grade glioma and glioblastoma combined). Vertical lines represent frequencies of various type of mutations at a given position. Synonymous, missense and truncating mutations are represented by green, blue and red lines, respectively. Gray lines are density curves. (**C**) Two-way clustering of driver genes and cancer types. Driver genes found in more than one cancer type are used (OGs in red and TSGs in blue). Abbreviations of cancer types are listed in Supplementary Table 2.

It is noteworthy that each of the 33 cancer types we examined involved at least one broad-spectrum driver and multiple tissue-specific drivers, implicating the synchrony of convergent and divergent disease pathways. Clustering of cancers based on broad-spectrum driver genes grouped cancer types largely matching their tissue and cellular origins (**Fig. 3C**).

#### Novel driver genes

We derived a list of high-confidence drivers consisting of 22 OGs and 74 TSGs with prediction specificities ≥0.99. By comparing this collection with known cancer drivers in the CGC annotations, we discovered two novel OGs and 28 novel TSGs. The two novel OGs, namely *CNOT9* in melanoma and *GTF2I* in thymoma, harbored single mutational hotspots disrupting highly conserved protein positions (**Fig. 4A-B**). The tumor-activating functions of these two genes have been confirmed by existing clinical and experimental studies (51–53). In particular, the *CNOT9* mutant is shown to stimulate stronger immune responses than the wild type, implicating the formation of a neoantigen that is a potential therapeutic target (52). The *GTF2I* mutant stimulates cell proliferation in vitro and has been associated with favorable prognosis of thymoma (51).

**Figure 4.**
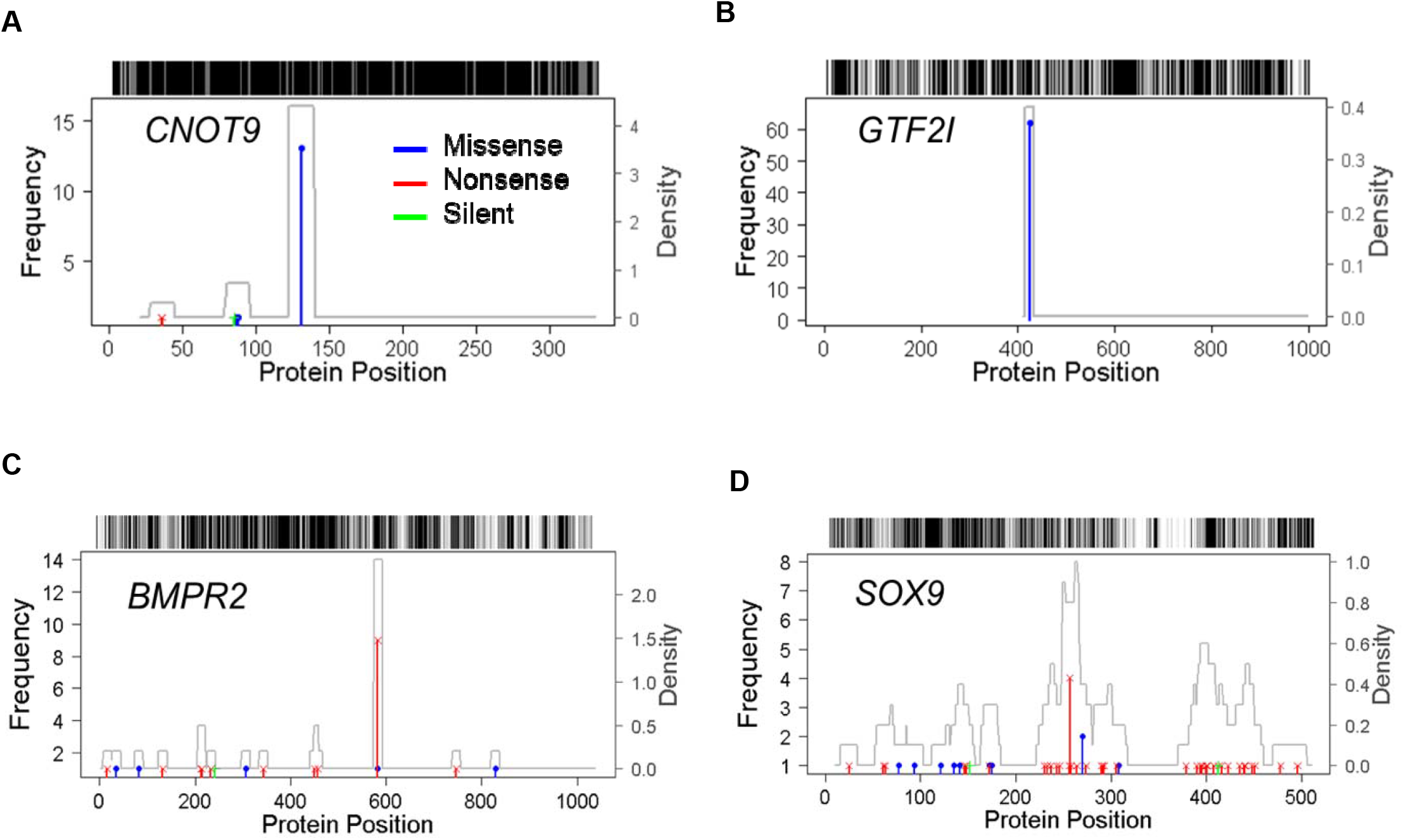
Mutational distributions of novel drivers. *CNOT9* (**A**) and *GTF2I* (**B**) are OGs. *BMPR2* (**C**) and *SOX9* (**D**) are TSGs.

All of the novel TSGs had an overabundance of truncating mutations (**Supplementary Fig. 1**). For example, frameshifting mutations in *SOX9* were observed in 40 colon cancers (**Fig. 4C**). As an atypical tumor suppressor, *SOX9* has been shown to interact with nuclear β-catenin. Inactivation of *SOX9* causes loss of inhibition of the oncogenic Wnt/β-catenin signalling pathway and is associated with patient survivals (54). Some novel TSGs harbor mutational hotspots. For instance, the N583 frameshifting mutation in *BMPR2* introduced premature stops of protein synthesis and was observed in nine stomach adenocarcinomas (**Fig. 4D**). All tumors carrying this mutation lost *BMPR2* expression, which in turn disrupted the TGF-β signalling pathway (55,56). Overall, our literature search found supporting evidence of tumor suppressing functions of 22 (78.6%) novel TSGs (**Supplementary Table 4**).

## DISCUSSION

Mounting evidence suggests tissue-specific organization of cancer pathways (3–5). Contextual classification of OGs, TSGs and PGs is a critical step to understanding the diverse molecular mechanisms and optimizing treatment plans. Although recurrence among patients has been taken as a surrogate of mutations under functional selection, recent investigations have shown that passenger hotspot mutations are common (57,58), which challenges approaches relying on ratiometric features. In this study, we addresses this issue by quantifying the contribution of genetic alterations to cancer fitness directly. The effectiveness of this approach can be demonstrated by the *MB21D2* gene with a C->T or C->G mutation at coding position 931 observed in multiple samples of various cancer types (**Fig. 5A-F**). Buisson et. al. showed that this mutational hotspot was due to its location in a hairpin loop susceptible to mutagenesis by *APOBEC3A*, thus functions as a passenger (57). GUST indeed estimated this gene was under neutral selection in individual cancer types and in combined samples (**Fig. 5G**) and classified it as a PG.

**Figure 5.**
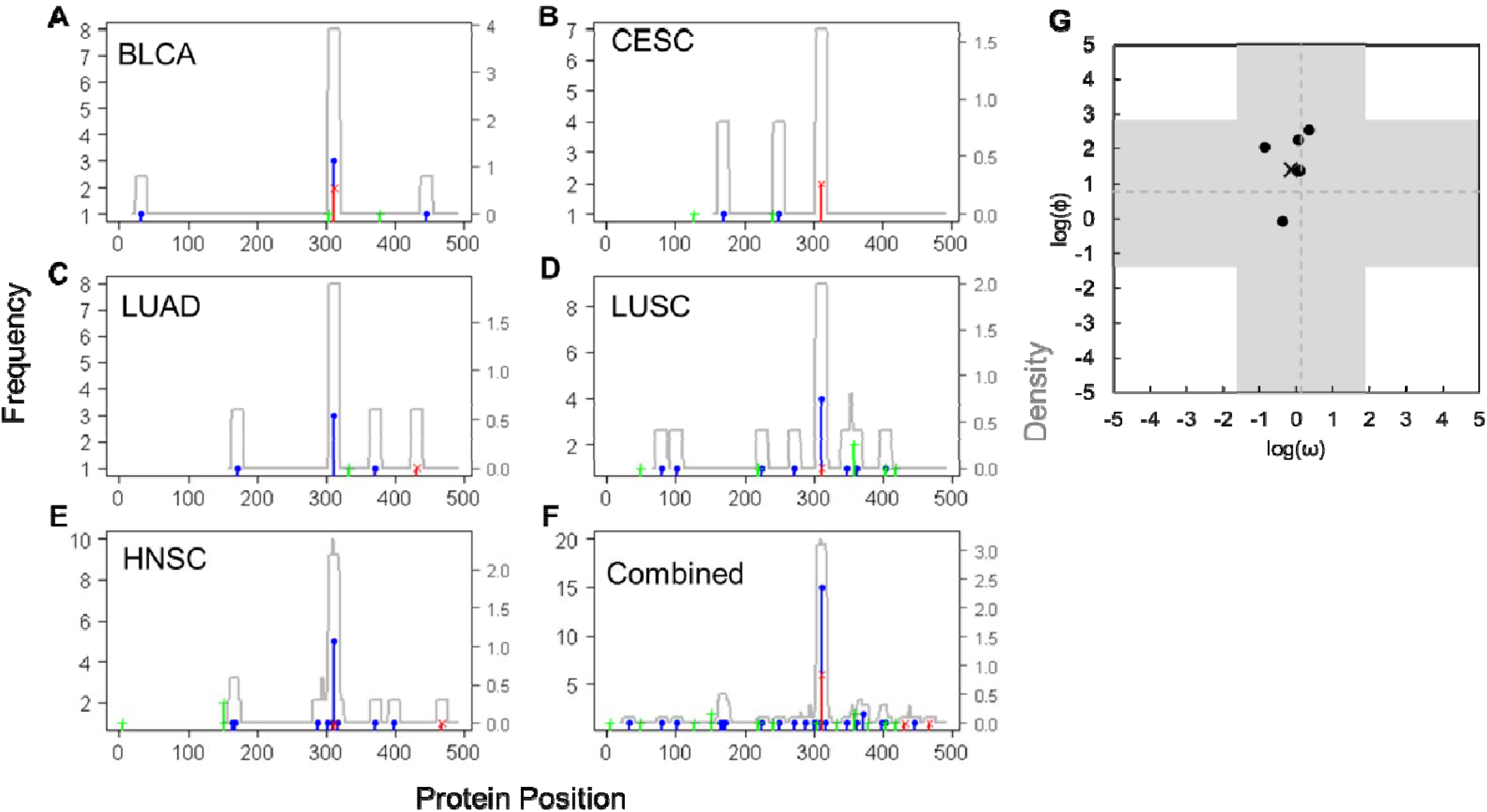
Mutational distributions and selection coefficients of the *MB21D2* gene. Distribution of mutations in bladder cancer (**A**), cervical cancer (**B**), head and neck cancer (**C**), lung adenocarcinoma (**D**), lung squamous cell carcinoma (**E**), and these cancer types combined (**F**). A mutation hotspot is located at protein position 311. (**G**) Selection coefficients estimated for individual cancer types (dots) and for combined samples (cross). Broken lines are the mean selection coefficient of all genes analyzed using all TCGA samples. Shaded areas are the 95% confidence intervals of the mean selection coefficients.

While GUST enables accurate predictions of driver genes, more information can be gathered by comparing features leading to these predictions across genes and across cancer types. As we have shown in our analysis, plotting the distributions of selection coefficients (**Fig. 1B**) and mutational hotspots (**Fig. 3B**) help interpret the classification results and establish connections to therapeutic targets and clinical outcomes. To facilitate these investigations, we have built an online database (https://compumedlab.net/gust/) with precomputed classification results from analyzing TCGA samples. Users can query the database and visually inspect somatic selection patterns and conservational patterns of selected genes. Combined with information showing if a gene has been annotated by CGC or OncoKB as a driver or a drug target, users can make informed decisions on prioritizing candidate genes for further investigations.

We compared the GUST method with the 20/20+ method, both of which employ a random forest model to integrate features extracted from somatic mutations. Despite that GUST uses only 10 features and 20/20+ uses 24 features, the accuracy of GUST is consistently higher. In the GUST model, selection coefficients contribute the most information content. The 20/20+ model does not use selection measures and the top four informative features are fractions of different types of mutations. These results suggest that using a small number of features engineered on biological premises is more powerful than feeding a large number of raw features to machine learning models. Furthermore, given the scarcity of known drivers for specific cancer types, reducing the number of features in predictive models helps mitigate overfitting problems.

Our analysis of the TCGA samples revealed contextual dependencies of cancer driver genes in several aspects. Consistent with previous reports, we found excessive drivers that promoted tumorigenesis in only one cancer type. The tissue specificities of broad-spectrum drivers are less obvious. We discovered that a majority of broad-spectrum OGs possessed multiple mutational hotspots affecting distinct functional domains. These hotspots are selectively engaged in various cancer types, suggesting divergent molecular mechanisms. Interestingly, we did not find any genes with dual OG/TSG roles. A straightforward explanation is that GUST makes predictions based on protein-altering substitutions and indels, thus unable to capture genes acting through other mechanisms, such as noncoding regulatory variants, copy number variants, differential expressions, post-translational modifications and epigenetic regulations. However, it also implicates that protein-altering mutations likely do not lead to opposite functional changes of the same gene. Further investigations of this hypothesis will shed light on key switches that divert paths of dual-role drivers.

As gene-centered treatment and drug repurposing attracts increasing interest (1,3,59), we expect the GUST method and the online database will facilitate discoveries of clinically actionable targets. The R implementation of the GUST algorithm is available on Github (https://github.com/liliulab/gust).

## DATA AVAILABILITY

Somatic mutations detected in TCGA samples are available from GDC data portal (https://portal.gdc.cancer.gov/).

Multiple sequence alignments of 100 vertebrate species are available from the UCSC Genome Browser (https://genome.ucsc.edu/).

20/20+ is an open source software available in the GitHub repository (https://github.com/KarchinLab/2020plus). Precomputed results of 20/20+ analysis of TCGA samples are available through the GDC data portal (https://gdc.cancer.gov/about-data/publications/pancan-driver).

## FUNDING

This work was supported by the National Institutes of Health [U54CA217376], the Flinn Foundation [#2088] and the ASU-Mayo Seed Grant.

## CONFLICT OF INTEREST

All authors claim no conflict of interest.

## SUPPLEMENTARY MATERIALS

**Supplementary Table 1**. Curated driver genes in various cancer types.

See the Excel file.

**Supplementary Table 2**. Number of driver genes predicted by GUST for each cancer type

See the Excel file.

**Supplementary Table 3**. Driver genes predicted by GUST from analyzing TCGA samples

See the Excel file.

**Supplementary Table 4**. Novel TSGs with supporting literature

See the Excel file.

**Supplementary Fig. 1.**
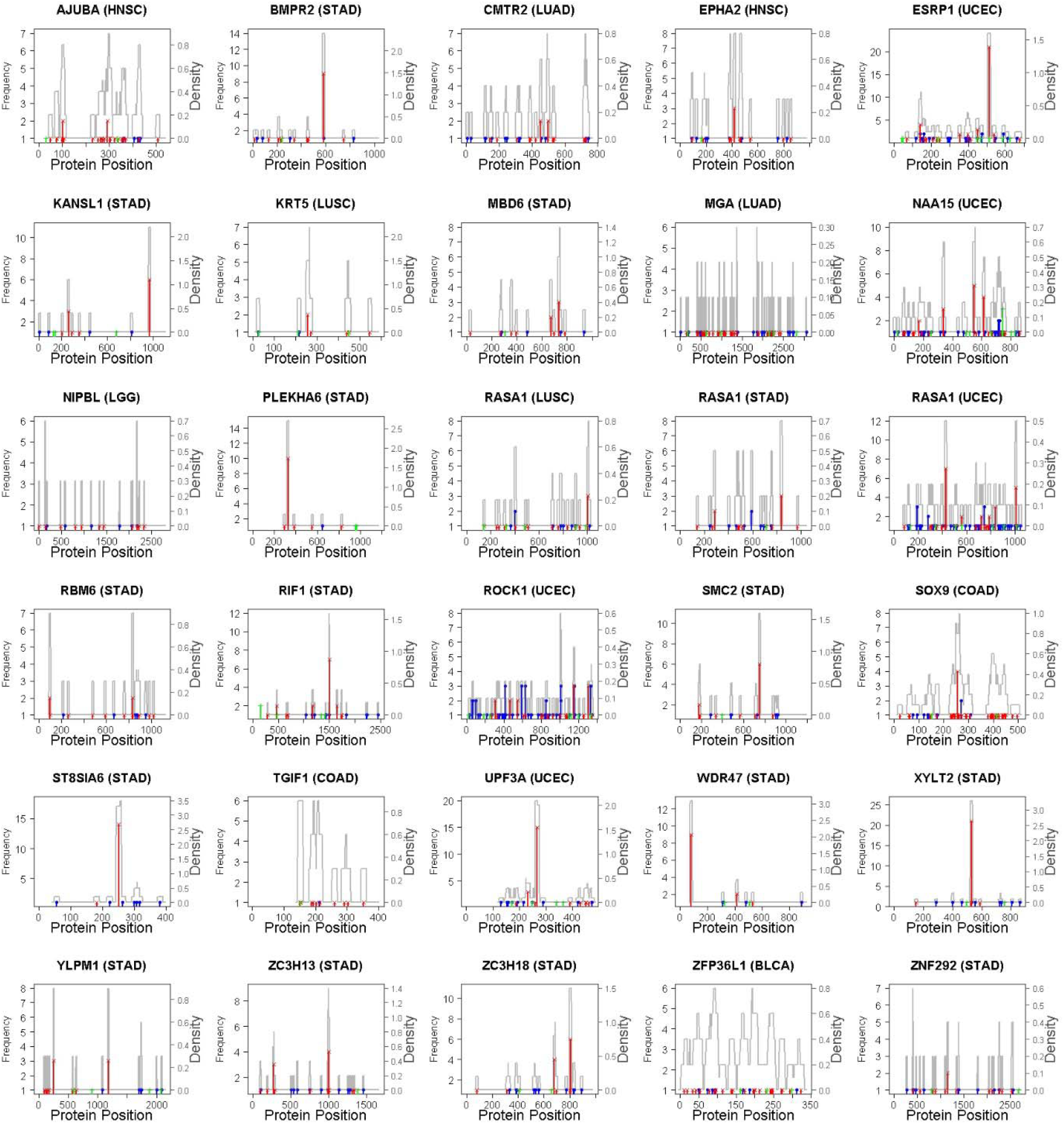
Mutational distribution of 28 novel TSGs.

